# PERFORATION-TYPE ANCHORS INSPIRED BY SKIN LIGAMENT FOR THE ROBOTIC FACE COVERED WITH LIVING SKIN

**DOI:** 10.1101/2024.01.24.576563

**Authors:** M. Kawai, M. Nie, H. Oda, S. Takeuchi

## Abstract

Skin equivalent, a living skin model composed of cells and extracellular matrix, possesses the potential as an ideal covering material for robots due to their biological functionalities. To employ skin equivalents as a coverage material for robots, a secure method for attaching them to the underlying structure is required. In this study, we developed a perforation-type anchor inspired by the structure of skin ligaments as a technique to effectively adhere skin equivalent to robotic surfaces, ensuring a smooth and non-protrusive integration, akin to natural skin adherence. We showed the effectiveness of plasma treatment for collagen-gel penetration within the perforation-type anchor. In characterizing the anchoring system, we investigated the impact of anchor diameter on preventing contraction and changes in holding strength. To showcase the versatility of perforation-type anchors in three-dimensional (3D) coverage applications, a 3D facial mold with intricate surface structure was covered with skin equivalent. Furthermore, we constructed a robotic face covered with dermis equivalent, capable of expressing smiles with the actuation to the skin equivalent through perforation-type anchors.

This research introduces an approach for the adhesion and actuation of skin equivalent with perforation-type anchors, potentially contributing to advancements in biohybrid robotics technology.

## INTRODUCTION

Robot skin materials serve multiple roles, particularly those for humanoid robots that require humanlike capabilities to operate in unpredictable and complex environments [1], [2]. Over time, a variety of robots mimicking these capabilities (e.g., tactile sensitivity [3], [4], self-repair [5], [6], perspiration [7], [8], and humanlike appearance [9]–[12]) have been developed. In contrast to existing robot skin materials, the use of cultured skin is expected to achieve the comprehensive biological functions of human skin [13] [14]–[16]. To apply cultured skin for robotics, it is necessary to construct 3D skin equivalents instead of the traditional two-dimensional (2D) tissue [14]–[17]. Currently, molding techniques that involve shrinking skin around an object have been employed to create 3D skin equivalents, which can evenly envelop 3D structures like robotic fingers [13] and grooves [18]. However, these methods have lacked a mechanism to fix the skin to the underlying subcutaneous layer, making the skin susceptible to significant deformation from external forces and raising the risk of damage. While protrusion anchors are commonly used as the traditional method for tissue fixation [13], [19]–[22], they pose drawbacks by protruding outward and being difficult to place on concave surfaces, which compromises the robot’s smooth appearance and design flexibility (refer to Figure S1).

In this paper, we describe a method employing “perforation-type anchors” (Figure 1A) inspired by skin ligaments. These ligaments, predominantly composed of collagen and elastin, are diminutive connective tissues that anchor the skin to the underlying tissues [19], enabling fluid facial expressions and bodily motions [20], [21]. By mimicking this structure, our method offers a less restrictive placement and maintains the robot’s aesthetic integrity. This involves applying a skin-forming cell-laden gel both onto and into specially designed V-shaped holes in the robot’s structure (Figure 1B).

**Fig. 1.**
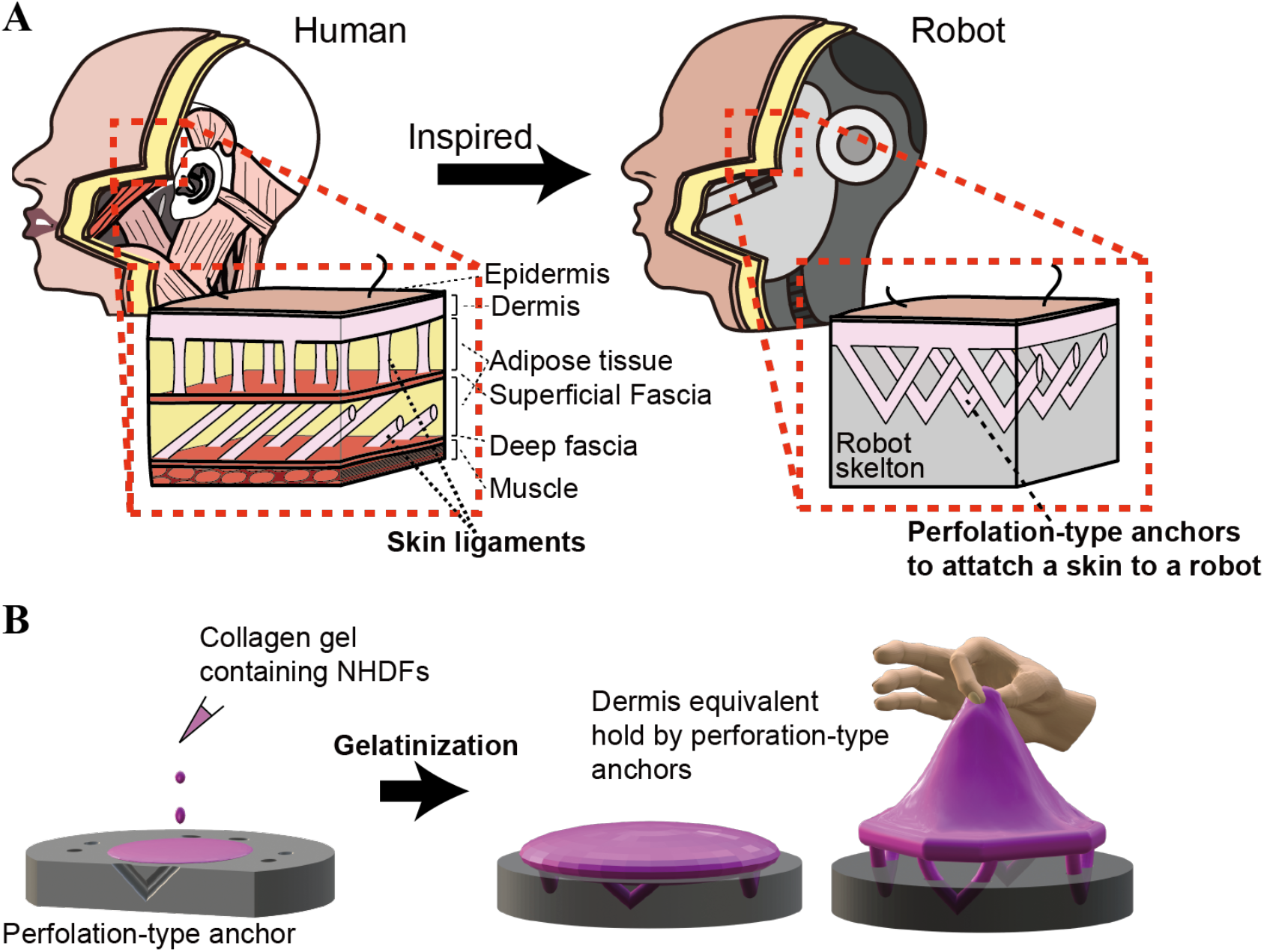
Conceptual illustration of the tissue-fixation method using perforation-type anchor. (**A**) Conceptual illustration of a perforation-type anchor inspired by skin ligaments to cover robots with skin equivalents (**B**) Process of the tissue fixation using perforation-type anchors.

We also introduce a water-vapor-based plasma treatment to enhance the gel’s penetration into these holes. Here, we establish a correlation between the anchor’s dimensions and its fixation efficacy via contraction and tensile testing. Furthermore, we demonstrate the versatility of this anchoring method through two distinct applications: the construction of 3D skin using a 3D facial mold covered with a skin equivalent, and the actuation of dermis on a robotic face designed to exhibit a smiling expression.

## RESULTS

### PLASMA TREATMENT FOR COLLAGEN PENETRATION WITHIN ANCHORS

Skin fixation using perforation-type anchors is achieved by gelating collagen gel introduced into the anchor’s interior. The anchoring process can fail when the collagen gel does not penetrate the perforation-type anchor’s interior or when air bubbles disrupt the internal tissue during the gelation. In cases where the anchors are not easily accessible, it becomes challenging to directly introduce the gel into the anchor’s interior, rendering the anchoring technique impractical. To address this issue, we demonstrated that the plasma treatment can make the device hydrophilic and enhance the gel penetration into the anchor’s interior. We initially evaluated the improvement in wettability by subjecting the surface of the 3D printed device to plasma treatment. Fig. 2A and 2B show a comparison of collagen contact angles on the device with and without plasma treatment, while Fig. 2C and 2D illustrate a comparison of the collagen spreading area. In samples without hydrophilization treatment, the average collagen contact angle was 37.9 degrees, and the collagen area was 18.0 mm^2^. In contrast, for samples subjected to plasma treatment, the collagen contact angle was reduced to 15.6 degrees, and the collagen area increased to 38.2 mm^2^. These results demonstrate that the plasma treatment employed in this study has successfully improved the wettability of the device.

**Fig. 2.**
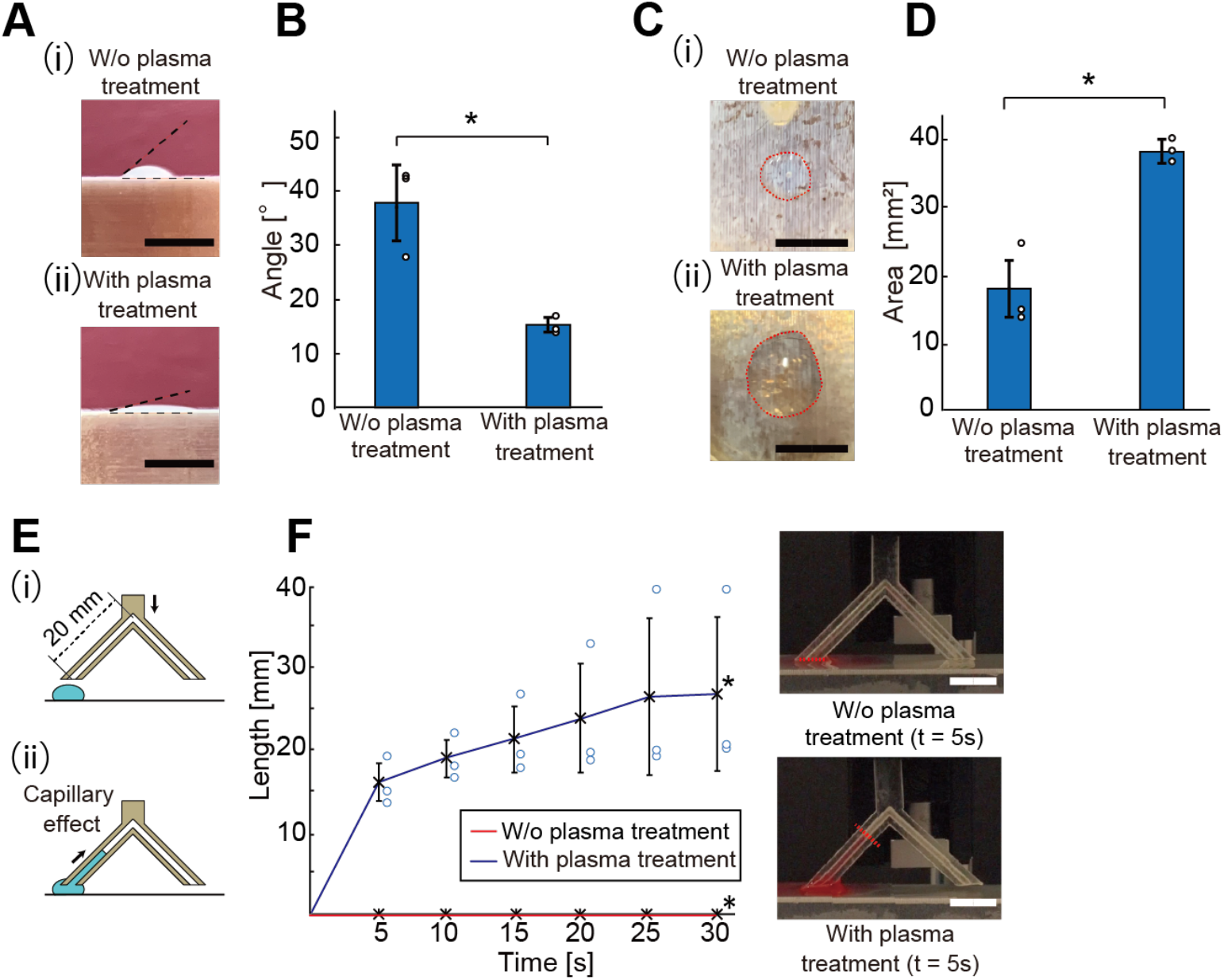
Validation of hydrophilic treatment effect on perforation-type anchors for enhanced collagen penetration. (**A**) Side-view images of collagen solution droplets on devices with and without plasma treatment. (**B**) Comparison graph of contact angles of collagen solution droplets on devices with and without plasma treatment (n = 3). The results are shown as the mean ± standard deviation (SD). *p < 0.05, unpaired Student’s t-test. (**C**) Top-view images of collagen solution droplets on devices with and without plasma treatment. (**D**) Comparison graph of collagen solution droplet area on devices with and without plasma treatment (n = 3). The results are shown as the mean ±SD. *p < 0.05, unpaired Student’s t-test. (**E**) Illustration of the experiment procedure to verify the hydrophilic treatment effect inside perforation-type anchors using capillary effect. (**F**) The results and images of the experiment to verify the plasma treatment effect inside perforation-type anchors using capillary effect (n = 3). The results are shown as the mean ±SD at each time point. *p < 0.05, unpaired Student’s t-test. Scale bars, (A) 10 mm; (C) 10 mm; (F) 10 mm.

To further validate the enhanced wettability within the anchor’s interior, we conducted a comparative experiment of the plasma treatment employing anchors with a diameter of 1 mm. Fig. 2E illustrates the experimental procedure. The device was equipped with a V-shaped anchoring hole with a diameter of 1.0 mm and was brought into contact with collagen. The subsequent changes in the liquid surface position due to capillary effect were observed. Fig. 2F and Movie S1 show the result of the experiment, comparing the rate at which collagen gel infiltrated the interior of the anchors. In the samples subjected to plasma treatment, the level of the collagen solution extended to a distance of 16.0 mm within 5 seconds of contact with the collagen solution. In contrast, for the samples without hydrophilization treatment, the collagen solution did not infiltrate the interior of the anchors at all. This observation demonstrates the effectiveness of plasma treatment in facilitating the entry of collagen solution into the anchor’s interior.

### EFFECTIVENESS OF PERFORATION-TYPE ANCHORS TO PREVENT THE CONTRACTION OF A DERMIS EQUIVALENT

To evaluate the ability of perforation-type anchors to suppress the contraction of skin equivalent, we performed a contraction test for the dermis equivalents with different perforation diameters. Fig. 3A shows the fabrication process of the samples for the contraction test. A collagen gel containing normal human dermal fibroblasts (NHDFs) was poured into a device with perforation-type anchors and gelated, resulting in the formation of a dermis equivalent. The dermis equivalent was cultured for seven subsequent days, during which the contraction process was observed.

**Fig. 3.**
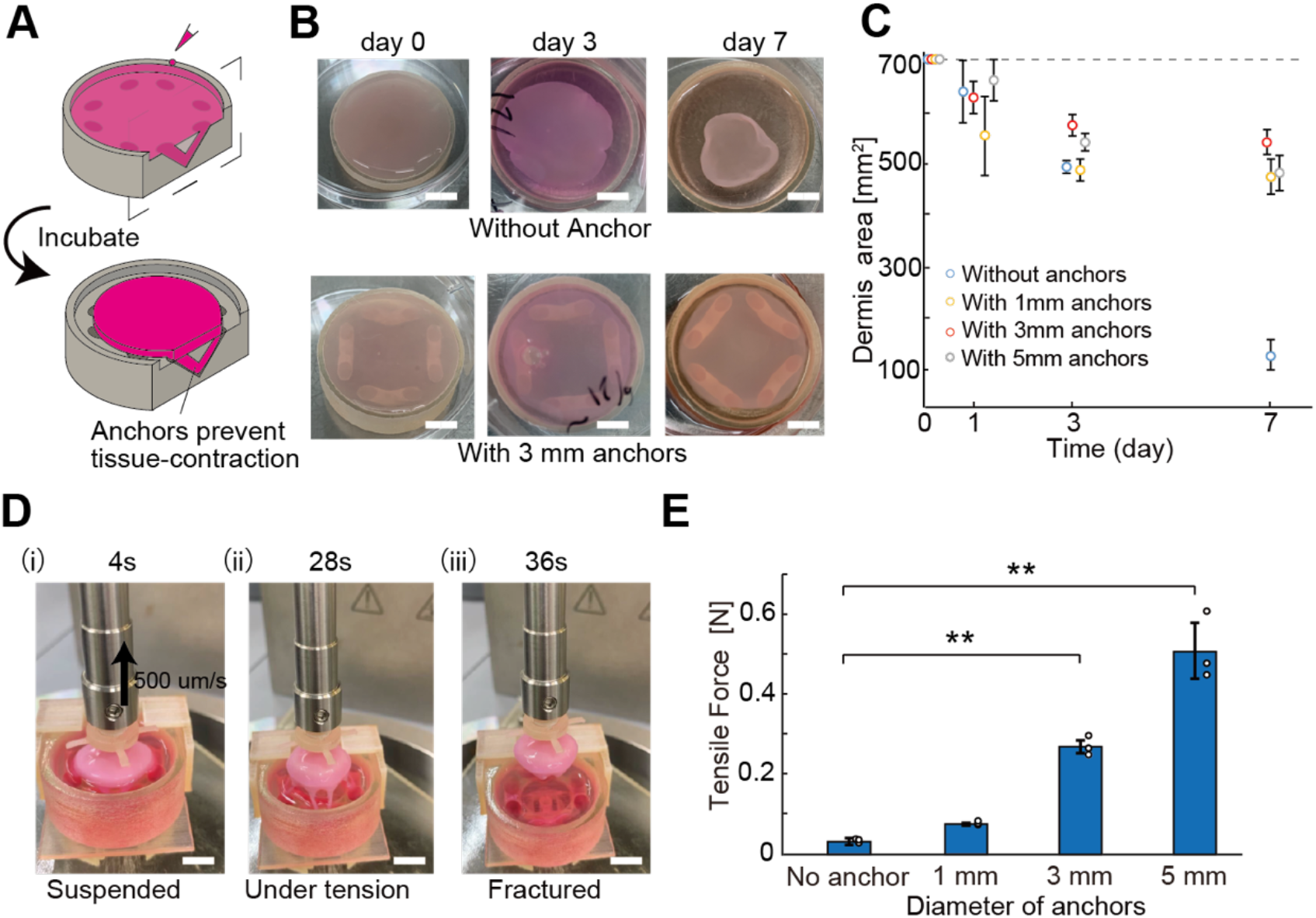
Functional evaluation of the perforation-type anchors to hold tissue. (**A**) Illustration of the device used to verify the effectiveness of perforation-type anchors on dermis equivalent. (**B**) Images of dermis contraction over time with and without 3 mm perforation-type anchors. (**C**) Comparison graph of dermis contraction over time with and without 3 mm perforation-type anchors (n = 3) The results are shown as the mean ± SD. (**D**) Images of the tensile test to confirm the retention strength of the perforation-type anchors (**E**) The result of the tensile test (n = 3). The results are shown as the mean ± SD. **p < 0.01, Tukey-Kramer test. Scale bars, (B) 10 mm; (D) 10 mm.

Fig. 3B shows comparative images of the samples without anchors and samples with 3 mm anchors immediately after gelation, after 3 days, and after 7 days. While the contraction of the tissue samples without anchors showed violent contraction, in the samples with 3 mm anchors, the contraction of the tissue samples with 3 mm perforation anchors was successfully restrained. Fig. 3C shows the graph comparing the area of the dermis equivalent after contraction. In devices without anchors, the dermis equivalent shrank up to 84.5% in 7 days, whereas with the anchored device, it was limited to 33.6% (for a diameter of 1 mm), 26.3% (for a diameter of 3 mm), and 32.2% (for a diameter of 5 mm). It can be inferred that even the perforation-type anchors with 1 mm diameter can prevent the shrinkage of dermis equivalent by holding tissue. The 3 mm anchors exhibit a higher resistance to contraction compared to the 1 mm anchors, but when the anchor diameter is increased to 5 mm, the post-contraction skin area is paradoxically reduced.

This phenomenon is likely attributable to the fact that the proportion of the area occupied by the anchors within the device surface becomes excessively large with 5 mm anchors; since the tissue shrinks towards its center, the inner peripheral line of the larger anchors is closer to the tissue center and results in larger shrinking distances. This result shows constraints on the practical implementation of perforation-type anchors with larger sizes.

### TENSILE TEST OF THE PERFORATION-TYPE ANCHOR

Tensile tests were performed to check the anchoring strength of the perforation-type anchors. Fig. S2 shows the setup to measure the anchoring strength between the dermis equivalent and the device. The testing setup consists of two devices: a device with anchors to hold a dermis sheet (dermis sheet holder) and a device for hooking and lifting the dermis-equivalent (hooking jig) (Figure S2(i)). The collagen containing NHDFs was poured and gelated on the dermis sheet holder with the hooking jig attached (Figure S2(ii-iii)). Following the 7-day incubation period after gelation, the tensile test was performed in which the hooking jig was pulled up with the dermis sheet fixed to the dermis sheet holder (Figure S2(iv)). The hooking jig was attached to a rheometer probe connected to motors and force sensors. This setup allowed for the pulling action while concurrently measuring the tensile force. Fig. 3D and Movie S2 show the images and movie of the tensile test. The dermis equivalent attached to the hooking jig was lifted, and the subsequent rupture of the anchor part was observed.

Fig. 3E illustrates a comparison graph of the maximum tensile strength across anchor diameter. In the devices without anchors, the maximum tensile strength was approximately 0.03 N, whereas in devices with anchors, it showed 0.08 N (for a diameter of 1 mm), 0.27 N (for a diameter of 3 mm), and 0.51 N (for a diameter of 5 mm), respectively. As the diameter of the anchor increases, a clear trend of increasing tensile strength can be observed. On the other hand, it’s important to note that there is a trade-off between the diameter and occupancy area of the perforation-type anchor. Larger anchor sizes impose stricter constraints on anchor placement. Designing anchors that align with the required fixed strength and placement constraints for a specific use case is essential.

### COVERAGE OF THE 3D FACIAL MOLD WITH A SKIN EQUIVALENT

To show the versatility of perforation-type anchors to cover 3D objects with intricate contours, we demonstrated the fabrication of the 3D facial device covered with a skin equivalent. Fig. S3 illustrates the process of constructing the skin equivalent covering the 3D facial device. This process involves a setup comprising three devices: the 3D facial device to be covered, an upper mold, and a lower mold (Figure S3(i)). The upper and lower mold create a space for filling and molding the collagen solution containing NHDFs (e.g. pre-gel dermis solution). The 3D facial device is equipped with eight perforation-type anchors on its sides for securing the skin equivalent. The fabrication process of the skin equivalent covering the 3D facial device involves two major steps: the construction of a dermis equivalent covering the 3D facial device (Figure S3(i-ii)) and the subsequent construction of the epidermis on the surface of the dermis equivalent (Figure S3(iii-v)).

The dermis equivalent covering the 3D facial device was created by pouring pre-gel dermis solution into the space in the assembled devices and incubating it for 7 days (Figure S3(ii)). During the incubation period, the pre-gel dermis solution is gelated while affixed to the anchors, resulting in the formation of the dermis equivalent fixed with the 3D facial device. Subsequently, the upper mold was removed from the 3D facial device (Figure S3(iii)) and epidermal tissue was formed on the dermis by seeding normal human epidermal keratinocytes (NHEKs) from the top of the dermis equivalent covering the 3D facial device. The tissue was cultured for 3 days in the medium and subsequent 14 days at the air-liquid interface (Figure S3(iv)). After the cultivation, the lower mold was removed, and the 3D facial device covered with skin equivalent was completed (Figure S3(v)).

Fig. 4 shows the images of the fabricated skin equivalent covering the 3D facial device during the fabrication process. Fig. 4A(i) shows the cultivation of the dermis equivalent covering the 3D facial mold, and Fig. 4A(ii) displays an image of the dermis equivalent after one week of cultivation. Fig. 4A(iii) represents the fabricated skin equivalent with dermis and epidermis layer, while Figure 4A(iv) shows the 3D facial device without the lower mold. The construction of the epidermis resulted in a change in the surface texture of the skin equivalent, giving it a whitish appearance. As shown in Fig. 4B(i), the cultured skin tissue is fixed to the device through perforation-type anchors. The skin equivalent is tightly secured to the device due to the tension during the contraction and fixation provided by the anchors, preventing it from detaching when pulled with force, as depicted in Fig. 4B(ii).

**Fig. 4.**
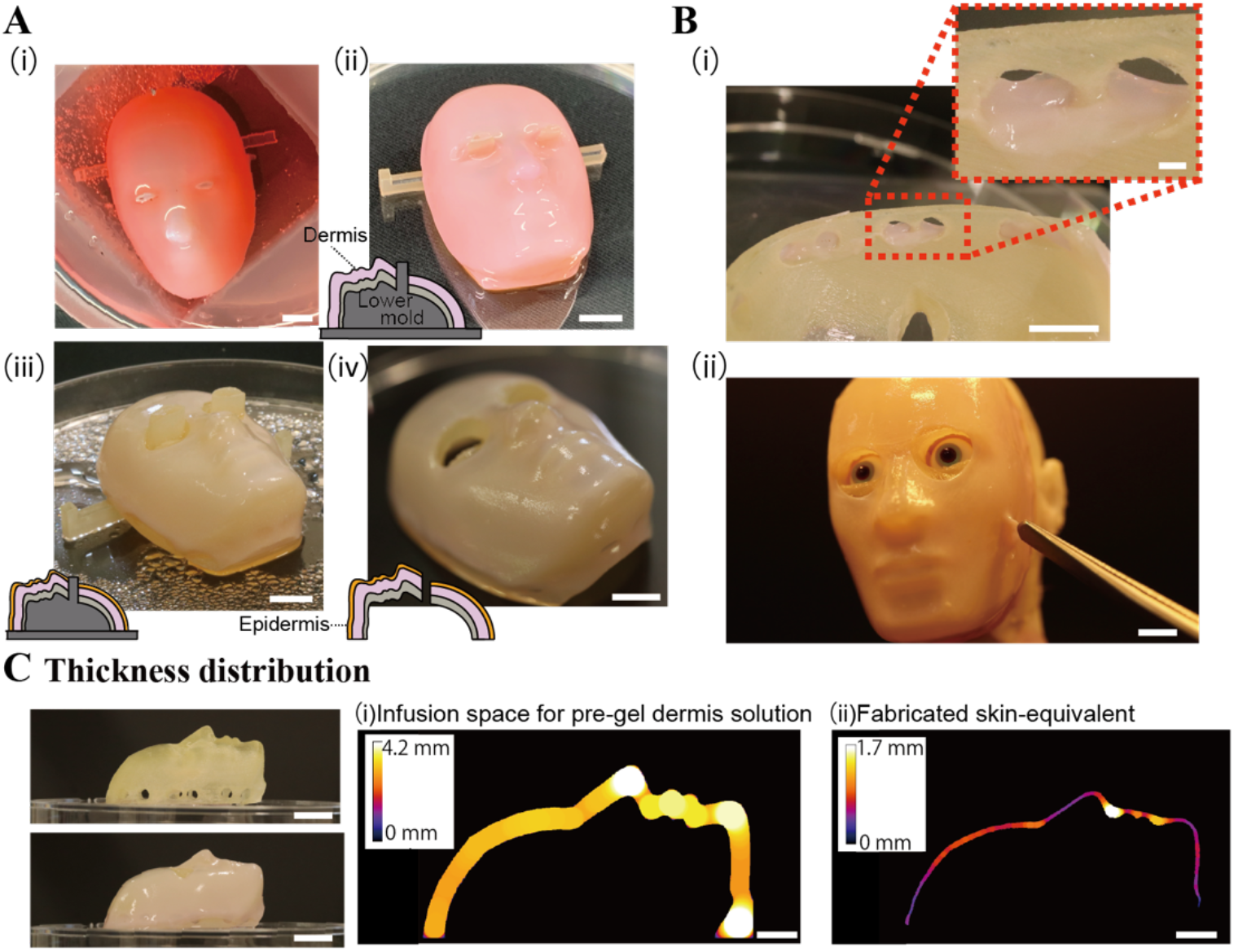
Demonstration of the perforation-type anchors to cover the facial device with skin equivalent. (**A**) Images of the fabrication process of a skin equivalent covering the 3D facial mold. (i-ii) The dermis equivalent anchored to the facial device was fabricated (iii-iv) Epidermis was formed on the surface of the dermis. (**B**) (i) An image of the back view of a facial device. Skin fixation through perforation-type anchors can be confirmed. (ii) The fixation through perforation-type anchors allows gently tugging on the skin. (**C**) Side view images of the facial device with and without a skin equivalent. Thickness maps of the infusion space for pre-gel dermis solution and the fabricated skin equivalent were measured. Scale bars, (A) 10 mm; (B) 10 mm, (inlet) 1 mm; (C) 10 mm, (thickness map) 5 mm.

Securing skin equivalent requires more than just friction; the use of perforation-type anchors is essential. Fig. S4 shows a skin equivalent created with a 3D facial device without perforation-type anchors as a control experiment. When anchors were not present, a skin equivalent was not fixed to the 3D facial device, and upon removing the upper mold, the tissue detached from the device (Figure S4A). Cultivating the tissue without fixation resulted in violent shrinkage, making it impossible to maintain the original shape (Figure S4B).

It is important to note that there is a limit to the contractile ability of skin equivalent, and our previous research has shown that achieving uniform coverage of 3D objects with skin equivalent also depends on the uniform thickness of the space into which collagen is injected [13]. Fig. 4C(i-ii) provides side-view photos of the facial device before and after its coverage by skin equivalent. Figure 4C(iii) illustrates the thickness distribution of the space where dermal solution is injected while figure 4C(iv) shows the thickness distribution of the skin equivalent after two weeks of epidermal seeding. The space within the device has an average thickness of 3.3 mm, while the resulting skin equivalent has an average thickness of around 0.81 mm. The standard deviation of the tissue thickness decreased from 0.50 mm immediately after gelation to 0.33 mm, demonstrating the ability to construct a skin equivalent with uniform thickness and a smooth surface. Upon closer examination, we observed slight variations in thickness in the constructed skin equivalent. Thinner skin tissue was observed in convex areas such as the top of the head, nose bridge, and chin, while relatively thicker skin formed in concave areas such as the border between the nose and forehead, below the nose, and around the mouth. These variations are due to differences in the direction of contraction forces caused by the contours. In convex areas, the contraction forces tend to result in thinner skin formation, while in concave areas, the opposite contraction forces may result in thicker skin formation. This fact can be a valuable aid in the design process when aiming for a more uniform skin construction.

### FABRICATION OF THE ROBOTIC FACE COVERED WITH DERMIS EQUIVALENT ACTUATED VIA PERFORATION-TYPE ANCHORS

In the human face, the enclosed mimetic muscles, like the zygomaticus major muscle, attach to the bone through retaining ligaments and to the skin through the skin ligament [20], [21]. In the expression of human emotions, particularly when smiling, the contraction of these mimetic muscles extending from the corners of the mouth to the cheeks lifts the corners of the mouth, creating elevated cheeks and inflamed subcutaneous tissue by muscle pump. Inspired by this smiling motion generated through the actuation of the zygomaticus major muscle, we created a robotic face covered with a dermis equivalent and a silicone layer connected to a slider via the perforation-type anchors. The sliding motion can produce a deformation of the silicone layer to generate the smiling expression (Figure 5A).

**Fig. 5.**
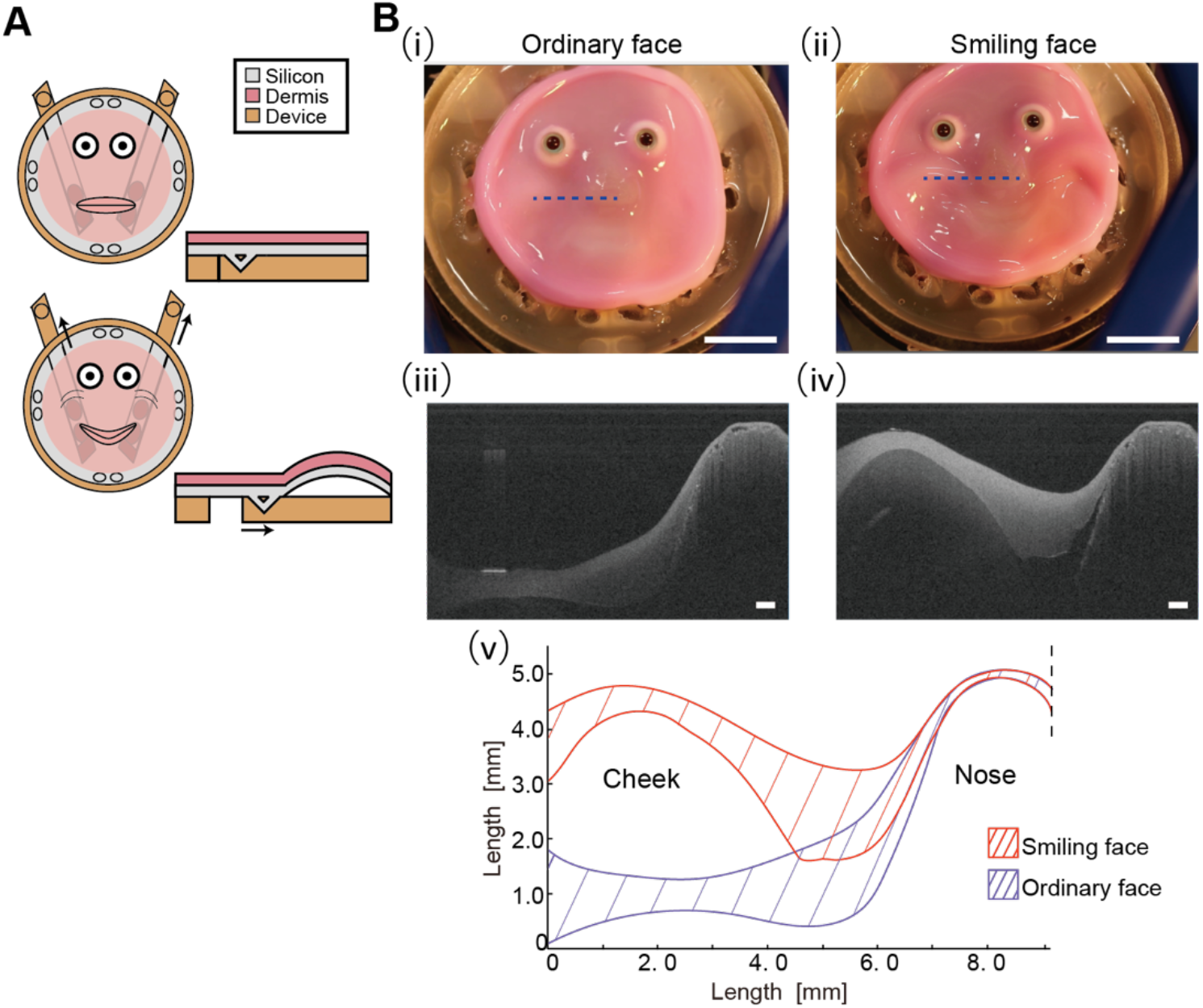
The smiling robotic face covered with dermis equivalent demonstrating actuation of the skin equivalent via the perforation-type anchors. (**A**) Conceptual illustration of the robotic face actuated with perforation-type anchors. (**B**) (i) Image of the robotic face with an ordinary expression. (ii) Image of the robotic face with a smiling expression. (iii) OCT image of the robotic face with an ordinary expression. (iv) OCT image of the robotic face with a smiling expression. (v) Comparison of the cross-sectional shape of skin during a smiling and an ordinary expression. Scale bars, (A) 10 mm; (B) 10 mm, 1 mm (OCT image).

Figure S5A illustrates the configuration of the robotic face covered with a dermis equivalent. The robotic face consists of a base part for dermis equivalent culture (base part) and two rod-shaped parts equipped with anchors to actuate the dermis equivalent (rod part). The base part features a silicone rubber layer, enabling the recreation of natural smiles by reproducing the bulging of subcutaneous tissue and muscles. As shown in Fig.S5B, the base part is equipped with 6 perforation-type anchors to fix the silicone layer and 7 perforation-type anchors to fix the dermis layer (Figure S5B). While anchors for the silicone layer and dermis equivalent are equipped with 3 mm perforation anchors, anchors for the rod parts are equipped with more secure 5 mm anchors, allowing them to pull the silicone layers more firmly.

Fig. 5B(i-ii) and Movie S3 display the smiling expression of the robotic face covered with a dermis equivalent. The silicon layer is pulled at the corners of the mouth by external mechanical actuators through perforation-type anchors, allowing the dermis equivalent at the corners of the mouth to be raised (Figure S6). The elevation of the subcutaneous tissue in the cheek area is also reproduced by the curvature of the pulled silicone layer, allowing the robotic face to display a natural smile. Fig. 5B(iii-iv) and Movie S4 is a cross-sectional image of the dermis equivalent from the cheek area to the nose area of the robotic face. Fig. 5B(v) shows a comparison graph of the cross-sectional shape of the dermis covering the robotic face during an ordinary expression and a smiling expression. These figures demonstrate that a smiling expression is created by selectively deforming the dermis equivalent of the cheek area via the perforation-type anchors. This result suggests the potential utility of perforation-type anchors for selective actuation of the skin on a face, like in-vivo facial muscles.

## DISCUSSION

Inspired by human skin ligaments, the current study designed a perforation-type anchor that enables the tissue fixation to the subcutaneous structure. This study highlights the hydrophilization of the device surface to address tissue infiltration into perforation-type anchors. This research explores the correlation between the size of perforation-type anchors and anchoring strength. As a demonstration of the tissue fixation, we applied perforation-type anchors to cover the 3D facial mold with a skin equivalent. Finally, a robotic face capable of generating smiling expressions via the perforation-type anchors was developed.

There remains an opportunity for exploring improved anchor shapes and placement. While a positive correlation has been identified between the size of perforation-type anchors and their holding capacity, a trade-off relationship exists between anchor size and the occupied area. The pursuit of an optimal anchor placement planning method stands as one of the next directions of this research. Additionally, there is untapped potential in exploring the more biomimetic shape of the perforation-type anchor. In the human body, skin ligaments are finer fibrous structures located within the subcutaneous tissue, maintaining a strong adhesion between the skin and underlying tissues [21]. The development of techniques to imitate this attachment with microscale mechanisms (e.g., the use of more intricate structures [22]–[24] or porous gels) may enable better adhesion, eliminating the need for punctures.

In this study, we explored the use of perforation-type anchors for applying a skin equivalent over complex 3D facial molds. Our previous research successfully produced a uniform thickness in skin equivalents covered on a robotic finger. This fabrication was achieved by designing a space for evenly pouring a pre-gel dermis solution. However, the facial shape, with its intricate unevenness, differs significantly from the simpler, convex shape of a finger. When applying the same pre-gel dermis solution technique on a 3D facial model, we noticed slight variations in skin thickness. This discrepancy may arise from the skin forming over concave and convex facial surfaces. It appears that the variation in skin contraction forces, which differ based on the surface curvature, impacts this outcome. On convex surfaces, these forces stretch the skin thinner, whereas on concave surfaces, they tend to lift the skin. Therefore, achieving precise skin thickness on facial models necessitates consideration of these differential forces.

The activity of facial muscles involved in forming expressions such as smiles is closely linked to the development of wrinkles [25], [26]. One significant next step in this research is to leverage this model to enhance our understanding of the mechanisms underlying wrinkle formation. Moreover, applying this knowledge to recreate such expressions on a chip could find applications in the cosmetics industry and the orthopedic surgery industry. Additionally, this study performed actuation on a dermis equivalent by controlling mechanical actuators positioned beneath the dermis equivalent. Substituting this mechanical actuator with cultured muscle tissue presents an intriguing prospect in the realization of a higher degree of biomimetics. Examining the correlation between facial muscle contractions and resulting facial expression can offer insights into the physiological aspects of emotion, leading to new exploration in the treatment of diseases, such as facial paralysis surgery.

## MATERIALS AND METHODS

### REAGANT

Dulbecco’s modified Eagle’s medium (DMEM) and penicillin-streptomycin were purchased from Sigma-Aldrich (St. Louis, MO, USA). Fetal bovine serum (FBS) was purchased from Biosera (Kansas City, MO, USA). L-ascorbic acid phosphate magnesium salt n-hydrate were purchased from Wako Pure Chemical Industries (Osaka, Japan). Keratinocyte Growth Medium 3 Kit was purchased from PromoCell GmbH (Heidelberg, Germany). 1x phosphate-buffered saline without Mg^2+^ Ca^2+^ (PBS(-)) was purchased from Cell Science & Technology Institute (Sendai, Japan). 10x phosphate-buffered saline without Mg^2+^ Ca^2+^ (10x PBS(-)) was purchased from Sigma-Aldrich (Japan). Type I collagen solution (IAC-50) was purchased from Koken Co., Ltd. (Tokyo, Japan). Ecoflex® (Supersoft 00–30) was purchased from Smooth-On, Inc. (Macungie, PA, USA).

### CELL CULTURE

Normal human dermal fibroblasts (NHDFs) were purchased from PromoCell GmbH and cultured in fibroblast growth medium (DMEM supplemented with 10% FBS, 1% penicillin-streptomycin, and 70 μg/mL L-ascorbic acid phosphate magnesium salt n-hydrate). Normal human epidermal keratinocytes (NHEKs) were purchased from PromoCell GmbH and cultured in keratinocyte growth medium (Keratinocyte Growth Medium 3 Kit supplemented with 1% penicillin-streptomycin). For both NHDFs and NHEKs, media were refurbished once every 2 days and the cells were subcultured at about 80% confluence. NHDFs were used within 10 passages and NHEKs were used within 8 passages. Fibroblast growth medium was used during the maturation of dermis-equivalents. Keratinocyte differentiation medium (1:1 mixture of fibroblast growth medium and keratinocyte growth medium) was used during and after the seeding of NHEKs onto dermis equivalents.

### FABRICATION OF THE SKIN EQUIVALENT

In this paper, the fabrication process of skin equivalents is primarily based on the methodology established by Bell et al [17]. The fabrication process of skin equivalents in this paper is divided into two procedures: fabrication of a dermis equivalent and construction of epidermis layer on the dermis equivalent. Dermis equivalent was fabricated by crosslinking the pre-gel dermis solution (a 9:1:5 mixed solution of type I collagen solution (IAC-50), 10× PBS(-), and fibroblast growth medium containing 1.2 × 10^6^ cells/ml NHDFs). At first, pre-gel dermis solution is poured into the device at room temperature (RT). Immediately after the pouring, the assembled device was placed in an incubator (37°C, 5% CO2) to crosslink the collagen, followed by culturing the tissue to allow for its maturation. During the maturation (7 days), cells adhered to the collagen matrix and remodeled it, leading to tissue contraction. In this paper, the contracted tissue (with 7 days of maturation culture) is referred to as ‘dermis equivalent’. Then, 200 μl of keratinocyte differentiation medium containing NHEKs at the concentration of 1.0 × 10^7^ cells/ml were seeded twice on the surface of the dermis equivalent with a one-hour interval. The dermis-equivalent (seeded with NHEKs) was cultured for 3 days in the keratinocyte differentiation medium and 2 weeks in the air-medium interface to allow the migration and differentiation of NHEKs. In this paper, the dermis equivalent covered with migrated and differentiated NHEKs is referred to as the ‘skin equivalent’.

### EVALUATION OF THE WATER VAPOR-BASED PLASMA TREATMENT

Plasma treatment was conducted using the water vapor-based plasma equipment (Aqua Plasma® Cleaner AQ-2000, SAMCO Inc.) [27]. The improvement of surface wettability on the device through plasma treatment was validated using the droplet method [28]. We prepared uniformly 3D-printed devices and coated them with parylene C. In this study, all 3D printed devices were printed by the 3D printer (AGILISTA-3100, KEYENCE Corp.) and coated with parylene C (2 μm thick) to prevent the release of cytotoxic substances from the devices. Then we divided the devices into two groups: an experimental group subjected to plasma treatment after parylene C coating and a control group left untreated with plasma treatment. Then, 10 μl of a collagen solution (a 9:1:5 mixed solution of type I collagen solution (IAC-50), 10× PBS(-), and fibroblast growth medium containing no NHDFs) was gently brought into contact with the surface of the device. The contact angle of the collagen droplets was calculated from side-view images using θ/2-method.

The verification of wettability improvement within the perforation-type anchor was conducted using the capillary effect. A device with a perforation-type anchor, with a diameter of 1 mm as shown in Fig. 2F, was prepared. After parylene C treatment, plasma treatment was applied to the experimental group. One end of the V-shaped anchor was then brought into contact with a collagen solution prepared on an aluminum plate placed on ice, as illustrated in Fig. 2F and Movie S1. The upward movement of the collagen surface owing to the capillary effect was recorded from the frontal perspective and compared with the negative control samples that did not undergo plasma treatment. In Fig. 2F and Movie S1, a small amount of red ink was used to stain the collagen gel for visualizing changes in the surface of the collagen solution.

### EVALUAITON OF THE EFFECTIVENESS OF PERFORATION-TYPE ANCHORS TO HOLD THE TISSUE

To investigate the function of perforation-type anchors in suppressing the contraction of skin equivalent, contraction tests were conducted on dermis equivalent cultured with devices with perforation-type anchors of different diameters. As depicted in Fig. 3A, a pre-gel dermis solution was poured into 3D-printed devices with perforation-type anchors and allowed to gel. All parts were printed by the 3D printer and conformally coated with parylene C. After gelation, the tissue was peeled off from the device surface to exclude adhesion between the device and the dermis equivalent. Subsequently, the tissue was cultured for 7 days, and photographs were obtained from the samples at intervals during this cultivation period. Image analysis using ImageJ was performed to observe changes in the area over the 7-day cultivation period.

### TENSILE TEST TO MEASURE THE ANCHORING STRENGTH OF THE PERFORATION-TYPE ANCHORS

To show the anchoring strength of the perforation-type anchors, tensile tests were performed as follows. The dermis equivalent was cultured using the devices as illustrated in Fig. S2. The device comprises two components: a dermis sheet holder and a hooking jig. The dermis sheet holder was equipped with perforation-type anchors of diameters 0 mm (without anchor), 1 mm, 3 mm, and 5 mm, respectively. pre-gel dermis solution was poured into the assembled device and incubated 7 days after the gelation. After the culture periods, the hooking jig was fitted to the shaft for disposable measuring systems (D-CP/PP) of a rheometer (MCR rheometer 302, Anton Paar Corp., Tokyo Japan) and the dermis sheet holder was secured onto a base fabricated using 3D printing. A tensile test was performed to measure the anchoring strength between the dermis-equivalent and dermis sheet holder by pulling the probe upward at a speed of 500 μm/s with tensile force recorded once every 0.25s.

### FABRICATION OF THE FACIAL DEVICE COVERED WITH SKIN EQUIVALENT

As shown in Fig. S2(i), the facial device consists of the upper mold, the facial device, and the lower mold, which were 3D-printed separately and assembled. All parts were printed by a 3D printer and conformally coated with parylene C. As shown in Fig. S2(ii), the pre-gel dermis solution, prepared on ice, was poured into the assembled device at room temperature (RT). Immediately after the pouring, the assembled device was placed in an incubator (37°C, 5% CO2) to crosslink the collagen and subsequently culture the tissue for maturation. 1 hour after the pouring, the upper mold was removed from the device to ensure an adequate oxygen supply to the tissue. During the maturation (7 days), cells adhered to and remodeled the collagen matrix, and the tissue contracts. Then, 200 μl of keratinocyte differentiation medium containing NHEKs at the concentration of 1.0 × 10^7^ cells/ml were seeded twice on the surface of the dermis equivalent with a one-hour interval (Figure S2(iii)). The dermis equivalent seeded with NHEKs was initially cultured for 3 days in the keratinocyte differentiation medium and for subsequent 14 days at the air-medium interface to allow the migration and differentiation of NHEKs. During the air-medium interface culture, the 3D facial device was cultured using a biological shaker to gently agitate the device and ensure that the skin covering the device did not dry out (Figure S2(iv)). After the cultivation period, the lower mold was pulled out from the device, and the skin equivalent covering the facial device was completed (Figure S2(v)).

### EVALUATION ON THE SHAPE OF THE 3D SKIN EQUIVALENTS

The thickness distribution of the fabricated skin equivalents was measured using Image J (Rasband, W.S., ImageJ, U. S. National Institutes of Health, Bethesda, Maryland, USA, https://imagej.nih.gov/ij/, 1997-2018). The 3D facial mold was captured from the side view before and after coverage with skin equivalent, positioned at the same location. The outlines of the skin equivalent in the lateral view were observed by taking the difference between the two images. Additionally, the thickness of the space intended for injection of pre-gel dermis tissue was calculated. The thickness distribution for each region was computed using ImageJ’s local thickness function [29]. Then, the average thickness and standard deviation were subsequently calculated.

### FABRICATION OF THE ROBOTIC FACE COVERED WITH DERMIS EQUIVALENT

As shown in Fig. S5, the robotic face consists of five parts and has the anchoring structure at the bottom part (i.e., part 5). The base part features six perforation-type anchors for the silicon layer fixture and the seven for dermis equivalent fixture. This silicone layer was incorporated to replicate the 3D puncture of subcutaneous tissue while smiling, contributing to the creation of a realistic smiling expression. This silicone layer was formed by cross-linking Ecoflex® within the base component, securely fastened to the anchors. To prevent Ecoflex® from infiltrating the anchors intended for fixing dermis equivalent, the cylindrical jigs were employed during the embedding process, ensuring the preservation of anchor locations. After the formation of the silicone rubber layer, the jigs were removed. Furthermore, individual silicon rubber-crafted noses and lips were seamlessly attached to the silicone rubber layers. a silicone layer was affixed to 5 mm perforation-type anchors in the rod parts, allowing for the actuation of the silicone layer. Following the construction of the silicone layer, a pre-gel dermis solution was poured into the base part. After gelation, the tissue was cultured for 7 days to construct the dermis equivalent covering the robotic face.

## Supplementary Materials

**Fig. S1**. Comparison between the conventional protrusion anchors and the proposed perforation-type anchors for tissue fixation.

**Fig. S2**. Preparation the tensile test device.

**Fig. S3**. The fabrication process of the 3D facial device covered with the skin equivalent

**Fig. S4**. The Dermis equivalent made on the 3D facial device without perforation-type anchors.

**Fig. S5**. Fabrication of the robotic face covered with dermis equivalent.

**Fig. S6**. The setup for the robotic face actuation.

**Movie S1**. The experiment to show the effectiveness of the plasma treatment.

**Movie S2**. The high-speed camera video of the tensile test.

**Movie S3**. The actuation of the robotic face covered with a dermis equivalent.

**Movie S4**. OCT microscopic imaging of the robotic face during smiling motion.

## Acknowledgments

The authors would like to thank SAMCO Inc. for technical support on the water vapor-based plasma equipment.

## Author contributions

Conceptualization: MK, NM

Methodology: MK

Investigation: MK, HO

Visualization: MK

Funding acquisition: ST

Project administration: NM, ST

Supervision: NM, ST

Writing – original draft: MK

Writing – review & editing: NM, HO, ST

## Data and materials availability

All data are available in the main text or the Supplementary Materials.

## References and Notes

[1] S. Kajita, H. Hirukawa, K. Harada, and K. Yokoi, “Introduction to Humanoid Robotics,” in Introduction to Humanoid Robotics, Berlin, Heidelberg: Springer Berlin Heidelberg, 2014, pp. 1–17. doi: 10.1007/978-3-642-54536-8_1.

[2] K. Hirai, M. Hirose, Y. Haikawa, and T. Takenaka, “The development of Honda humanoid robot,” in Proceedings. 1998 IEEE International Conference on Robotics and Automation (Cat. No.98CH36146), Leuven, Belgium: IEEE, 1998, pp. 1321–1326. doi: 10.1109/ROBOT.1998.677288.

[3] L. E. Osborn et al., “Prosthesis with neuromorphic multilayered e-dermis perceives touch and pain,” Sci. Robot., vol. 3, no. 19, p. eaat3818, Jun. 2018, doi: 10.1126/scirobotics.aat3818.

[4] Y. Yan et al., “Soft magnetic skin for super-resolution tactile sensing with force self-decoupling,” Sci. Robot., vol. 6, no. 51, p. eabc8801, Feb. 2021, doi: 10.1126/scirobotics.abc8801.

[5] S. Terryn, J. Brancart, D. Lefeber, G. Van Assche, and B. Vanderborght, “Self-healing soft pneumatic robots,” Sci. Robot., vol. 2, no. 9, p. eaan4268, Aug. 2017, doi: 10.1126/scirobotics.aan4268.

[6] C. B. Cooper et al., “Autonomous alignment and healing in multilayer soft electronics using immiscible dynamic polymers,” Science, vol. 380, no. 6648, pp. 935–941, 2023, doi: 10.1126/science.adh0619.

[7] Y. Asano et al., “Human mimetic musculoskeletal humanoid Kengoro toward real world physically interactive actions,” in 2016 IEEE-RAS 16th International Conference on Humanoid Robots (Humanoids), Cancun, Mexico: IEEE, Nov. 2016, pp. 876–883. doi: 10.1109/HUMANOIDS.2016.7803376.

[8] A. K. Mishra et al., “Autonomic perspiration in 3D-printed hydrogel actuators,” Sci. Robot., vol. 5, no. 38, p. eaaz3918, Jan. 2020, doi: 10.1126/scirobotics.aaz3918.

[9] J. Oh, D. Hanson, W. Kim, Y. Han, J. Kim, and I. Park, “Design of Android type Humanoid Robot Albert HUBO,” in 2006 IEEE/RSJ International Conference on Intelligent Robots and Systems, Beijing, China: IEEE, Oct. 2006, pp. 1428–1433. doi: 10.1109/IROS.2006.281935.

[10] C. Becker-Asano and H. Ishiguro, “Evaluating facial displays of emotion for the android robot Geminoid F,” in 2011 IEEE Workshop on Affective Computational Intelligence (WACI), Paris, France: IEEE, Apr. 2011, pp. 1–8. doi: 10.1109/WACI.2011.5953147.

[11] J. Parviainen and M. Coeckelbergh, “The political choreography of the Sophia robot: beyond robot rights and citizenship to political performances for the social robotics market,” AI Soc., vol. 36, no. 3, pp. 715–724, Sep. 2021, doi: 10.1007/s00146-020-01104-w.

[12] N. Mori, Y. Morimoto, and S. Takeuchi, “Skin integrated with perfusable vascular channels on a chip,” Biomaterials, vol. 116, pp. 48–56, Feb. 2017, doi: 10.1016/j.biomaterials.2016.11.031.

[13] M. Kawai, M. Nie, H. Oda, Y. Morimoto, and S. Takeuchi, “Living skin on a robot,” Matter, vol. 5, no. 7, pp. 2190–2208, Jul. 2022, doi: 10.1016/j.matt.2022.05.019.

[14] Y. Xie et al., “Development of a Three-Dimensional Human Skin Equivalent Wound Model for Investigating Novel Wound Healing Therapies,” Tissue Eng. Part C Methods, vol. 16, no. 5, pp. 1111–1123, Oct. 2010, doi: 10.1089/ten.tec.2009.0725.

[15] S. Inagaki, Y. Morimoto, I. K. Suzuki, K. Emoto, and S. Takeuchi, “Co-culture system of human skin equivalents with mouse neural spheroids,” J. Biosci. Bioeng., vol. 136, no. 3, pp. 239–245, 2023, doi: 10.1016/j.jbiosc.2023.05.008.

[16] J. Lee et al., “Hair-bearing human skin generated entirely from pluripotent stem cells,” Nature, vol. 582, no. 7812, pp. 399–404, Jun. 2020, doi: 10.1038/s41586-020-2352-3.

[17] E. Bell, H. P. Ehrlich, D. J. Buttle, and T. Nakatsuji, “Living Tissue Formed in Vitro and Accepted as Skin-Equivalent Tissue of Full Thickness,” Science, vol. 211, no. 4486, pp. 1052–1054, 1981, doi: 10.1126/science.7008197.

[18] A. Pappalardo et al., “Engineering edgeless human skin with enhanced biomechanical properties,” Sci. Adv., vol. 9, no. 4, p. eade2514, Jan. 2023, doi: 10.1126/sciadv.ade2514.

[19] C. Stecco, Functional atlas of the human fascial system. Elsevier Health Sciences, 2014.

[20] B. C. Mendelson and S. R. Jacobson, “Surgical Anatomy of the Midcheek: Facial Layers, Spaces, and the Midcheek Segments,” Clin. Plast. Surg., vol. 35, no. 3, pp. 395–404, Jul. 2008, doi: 10.1016/j.cps.2008.02.003.

[21] N. Corduff, “Neuromodulating the SMAS for Natural Dynamic Results,” Plast. Reconstr. Surg. - Glob. Open, vol. 9, no. 8, p. e3755, Aug. 2021, doi: 10.1097/GOX.0000000000003755.

[22] G. Ma and C. Wu, “Microneedle, bio-microneedle and bio-inspired microneedle: A review,” J. Controlled Release, vol. 251, pp. 11–23, Apr. 2017, doi: 10.1016/j.jconrel.2017.02.011.

[23] D. Han et al., “4D Printing of a Bioinspired Microneedle Array with Backward-Facing Barbs for Enhanced Tissue Adhesion,” Adv. Funct. Mater., vol. 30, no. 11, p. 1909197, Mar. 2020, doi: 10.1002/adfm.201909197.

[24] K.-Y. Seong et al., “A self-adherent, bullet-shaped microneedle patch for controlled transdermal delivery of insulin,” J. Controlled Release, vol. 265, pp. 48–56, Nov. 2017, doi: 10.1016/j.jconrel.2017.03.041.

[25] N. Puizina-Ivić, “Skin aging.,” Acta Dermatovenerol. Alp. Pannonica Adriat., vol. 17, no. 2, pp. 47–54, Jun. 2008.

[26] J. Genzer and J. Groenewold, “Soft matter with hard skin: From skin wrinkles to templating and material characterization,” Soft Matter, vol. 2, no. 4, pp. 310–323, 2006, doi: 10.1039/B516741H.

[27] H. Terai, R. Funahashi, T. Hashimoto, and M. Kakuta, “Heterogeneous bonding between cyclo-olefin polymer (COP) and glass-like substrate by newly developed water vapor-assisted plasma, Aqua Plasma Cleaner,” Electr. Eng. Jpn., vol. 205, no. 4, pp. 48–56, Dec. 2018, doi: 10.1002/eej.23164.

[28] T. Young, “III. An essay on the cohesion of fluids,” Philos. Trans. R. Soc. Lond., vol. 95, pp. 65–87, 1805, doi: 10.1098/rstl.1805.0005.

[29] T. Hildebrand and P. Rüegsegger, “A new method for the model-independent assessment of thickness in three-dimensional images,” J. Microsc., vol. 185, no. 1, pp. 67–75, Jan. 1997, doi: 10.1046/j.1365-2818.1997.1340694.x.

