## Supplemental figures for "PERFORATION-TYPE ANCHORS INSPIRED BY SKIN LIGAMENT FOR THE ROBOTIC FACE COVERED WITH LIVING SKIN"

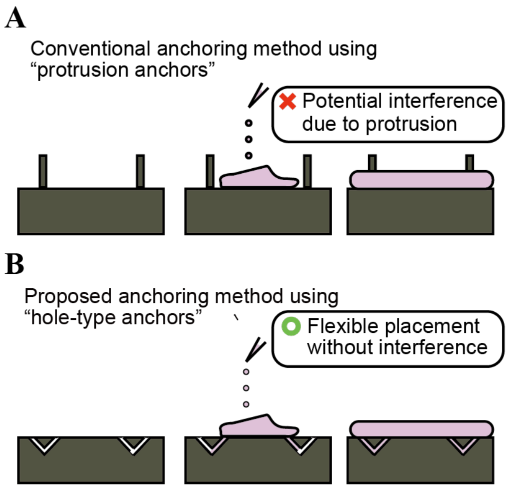


**Fig. S1. Comparison between the conventional protrusion anchors and the proposed perforation-type anchors for tissue fixation.** (A) Mechanism of the protrusion anchors for securing tissues. (B) Mechanism of the perforation anchors for securing tissues.

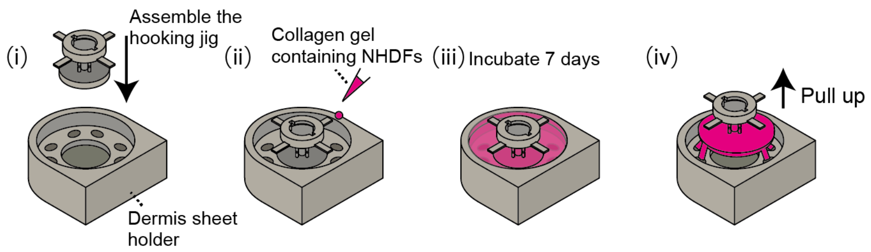


**Fig. S2. Preparation of the tensile test setup.**

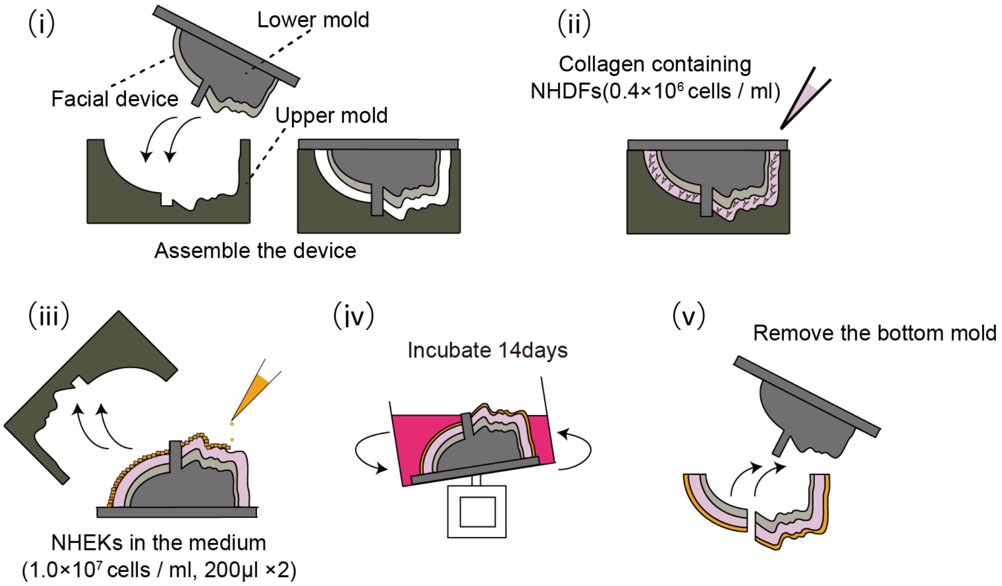


**Fig. S3. The fabrication process of the 3D facial device covered with the skin equivalent** (i-ii) The dermis equivalent anchored to the 3D facial device was fabricated (iii-iv) epidermis was formed on the surface of the dermis equivalent.

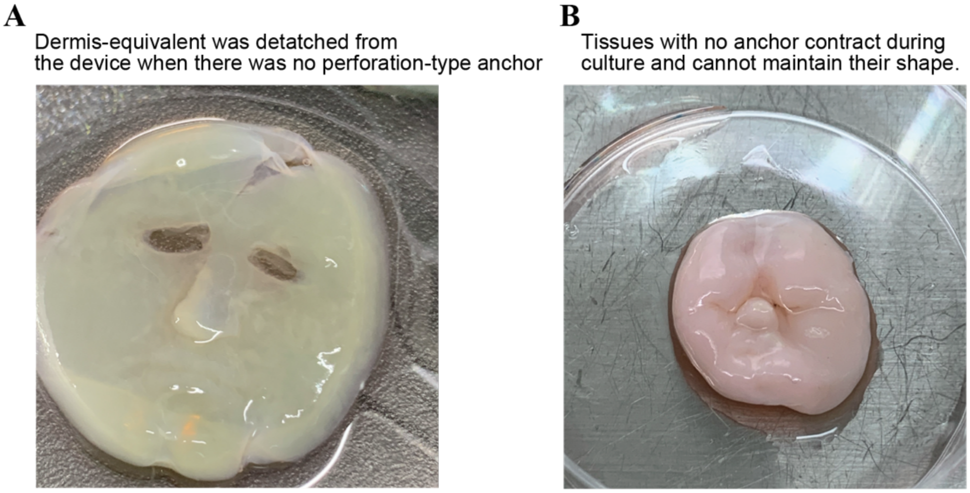


**Fig. S4. The Dermis equivalent made on the 3D facial device without perforation-type anchors.** (**A**) In the absence of perforation-type anchors, the dermis equivalent detached from the facial device when peeling off the upper mold. (**B**) Cultivating dermis equivalent that are not secured by perforation-type anchors results in an inability to maintain their shape.

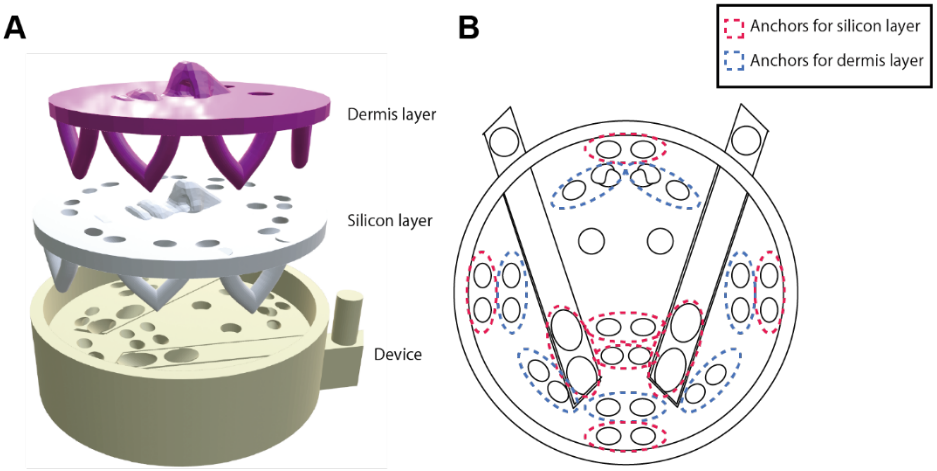


**Fig. S5. Fabrication of the robotic face covered with dermis equivalent.** (**A**) Composition of the robotic face covered with dermis equivalent (**B**) Number and arrangement of perforation-type anchors in the base part.

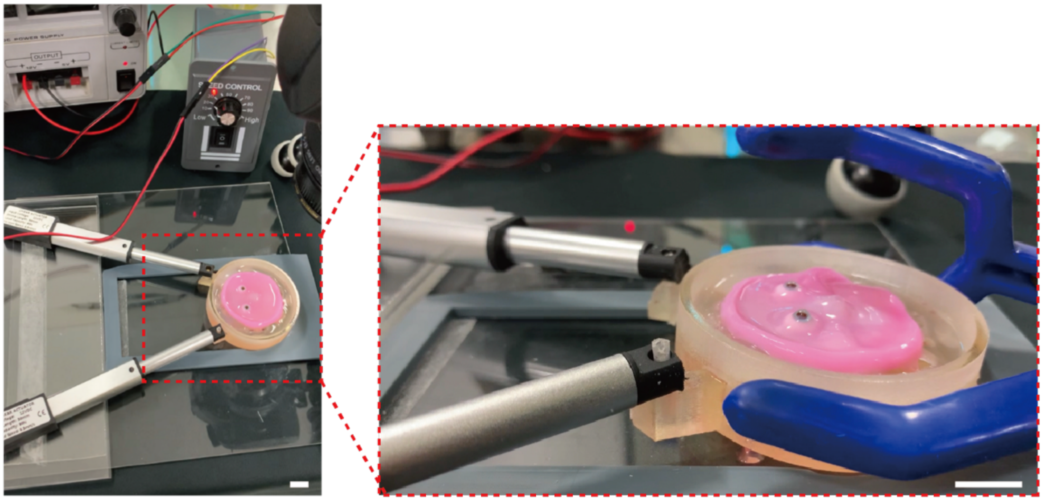

**Fig. S6. The Setup for the robotic face actuation. Scale bar, 10 mm.**
